# A high-quality genome resource for *Cercospora* cf. *flagellaris*, a causal agent of Cercospora leaf blight of soybeans

**DOI:** 10.64898/2026.06.16.732711

**Authors:** Zachary A. Carver, Trey Price, Jonathan K. Richards, Vinson P. Doyle

## Abstract

A highly contiguous and complete reference genome of *Cercospora* cf. *flagellaris*, the causal agent of foliar disease on many plant hosts including Cercospora leaf blight of soybean, was assembled using a combination of PacBio and Illumina sequencing reads. The genome assembly is 33.72 Mb in length and consists of 14 nuclear scaffolds and one mitochondrial contig. Four scaffolds have telomeric repeats on both ends and represent fully assembled chromosomes, while nine scaffolds represent partially assembled chromosomes with telomeric repeats on one end. The assembly has an N50 of 2.90 Mb and an L50 of 5 scaffolds. Genome annotation identified 11,268 genes, of which 947 and 360 were predicted to encode secreted proteins and effectors, respectively. Additionally, 512 genes were predicted to encode carbohydrate-active enzymes and 60 biosynthetic gene clusters were annotated. Taken together, this annotated genome assembly will be a valuable resource for genomics, host-pathogen interactions, and population biology research in this economically important pathosystem.

## Resource Announcement

Cercospora leaf blight (CLB) is a highly destructive soybean disease. In the southern U.S., yield losses attributed to CLB in 2025 were estimated at over 4.3 million bushels (Crop Protection Network 2026). Historically, CLB was solely attributed to the causal agent *C. kikuchii*, however, recent studies have demonstrated that multiple *Cercospora* species can cause CLB and that the predominant species can vary among geographical regions (Albu et al. 2016). In the United States, *C*. cf. *flagellaris* is the predominant species associated with CLB, although *C*.*sigesbeckiae, C. kikuchii, C. iranica*, and other species have been associated with the disease (Albu et al. 2016; Shrestha et al. 2024). Besides soybean, *C*. cf. *flagellaris* can infect a moderately broad range of hosts including pokeweed (Ellis and Martin 1882), industrial hemp (Doyle et al. 2019), cotton (Albu et al. 2016), and okra (Chai et al. 2021). Despite its economic importance, limited genomics resources exist for *C*. cf. *flagellaris* which hinders fundamental research on host-pathogen interactions, population biology, and host adaptation within this pathosystem. To address this knowledge gap, we generated the first annotated reference genome of *C*. cf. *flagellaris* using a combination of long and short read sequencing.

The *C*. cf. *flagellaris* isolate DMCC1359 was isolated from a symptomatic leaf of *Triodanis perfoliata* on the edge of a soybean field at the Ben Hur Research Station (30.376688, - 91.16874) in Baton Rouge, LA on August 16^th^, 2016, during a survey for alternative hosts for *Cercospora*. Surveys from potential alternative hosts as well as soybean plants exhibiting symptoms of Cercospora leaf blight took place from June to September in 2016 and 2017 when the incidence of leaf pathogens was highest and co-incident with the presence of CLB in soybean fields. All symptomatic foliar tissue were transported to LSU, Baton Rouge, LA in Zip-loc bags on ice. Foliar tissue was examined beneath a stereoscope for fasciculate cercosporoid conidiophores. Leaves without detectable conidiogenous structures were surface sterilized for 30s in 70% ethanol and then in 10% bleach, rinsed three times with DI, patted dry with sterile paper towels, and incubated at RT in high relative humidity for no more than 72hr before re-examination. A flamed steel inoculating needle dipped in sterile 15% glycerol was used to obtain and transfer single conidium or single conidiophore isolates to fresh potato dextrose agar (PDA) amended with streptomycin (30 μg/L). Plates were incubated at 25°C in darkness for 7d or until cultures were approximately 50 mm in diameter prior to DNA extraction. DNA was extracted from mycelia scraped from colonies on PDA using an amended cetyl trimethylammonium bromide (CTAB) DNA extraction protocol (Doyle and Doyle 1990). High-quality DNA was quantified by NanoDrop™ spectrophotometer (Thermo Fisher Scientific, Waltham, MA, U.S.A.) and diluted to a working concentration of 25 ng/μL. We next used PCR to amplify five markers commonly used in *Cercospora* species identification (calmodulin, actin, histone3, nrITS, and tef1-α), as in Albu et al. 2016, to identify the isolates we collected from alternative hosts.

Multilocus sequence alignments of representative isolates from this study and data from previous phylogenetic studies of *Cercospora* (Albu et al. 2016; Bakhshi et al. 2015; Groenewald et al. 2013; Soares et al. 2015) were estimated using MAFFT v7.310 (Katoh et al. 2017; Kuraku et al. 2013) under G-INS-i iterative refinement strategy, a 200PAM/k=2 scoring matrix for nucleotide sequences, 1.53 gap opening penalty, and a 0.0 offset value and trimmed in AliView version 1.18.1. Species assignments were made based on the multilocus maximum likelihood phylogeny inferred in RaXML (raxmlHPC-HYBRID) applying a general time reversible model of nucleotide evolution and gamma distribution on rate variation across sites (including invariant sites) coupled with rapid bootstrapping (Stamatakis 2014). Seventeen isolates of *Cercospora* from alternative hosts were identified based on the multilocus phylogenetic analysis. Seven isolates were recovered from *Phytolacca americana* (pokeweed), two from *Sorghum halepense* (Johnsongrass), one from a *Cyperus esculentus* (Yellow nutsedge), three from *Gossypium hirsutum* (cotton), one from *Morus alba* (mulberry), one from *Ambrosia trifida* (giant ragweed), one from *Cornus florida* (dogwood), and one from *Triodanis perfoliata* (clasping Venus’ looking glass). These isolates were assigned to *Cercospora sp*. A (nomenclature adopted from Groenewald et al. 2013), *C*. cf. *sigesbeckiae*, and *C*. cf. *flagellaris. Cercospora sp*. A was found on *P. americana* (N=2) and *Sorghum halepense* (N=2). Two isolates of *C*. cf. *sigesbeckiae* were isolated from *G. hirsutum*. Five isolates of *C*. cf. *flagellaris* were recovered from *P. americana*, one from a *C. esculentus*, one from *G. hirsutum*, one from *C. florida*, one from *A. trifida*, one from *T. perfoliata*, and one from *M. alba* (Figure 1, Supplementary Table 1). Gene sequences were deposited into the NCBI database (Supplementary Table 1). Isolate DMCC1359 was confirmed as *C*. cf. *flagellaris* and selected for reference genome development. This isolate was selected for sequencing because *Cercospora speculariae* was described in 1888 by Ellis and Langlois from a specimen collected from *Triodanis perfoliata* (syn. *Specularia perfoliata*) in Louisiana, and at the time of Chupp’s monograph on the genus Cercospora it was only known from the type locality (Chupp 1954). Figure 1 suggests *C. speculariae* and *C. flagellaris* may be synonyms, but examination of the type specimens for each need to be consulted in order to make a final determination.

**Figure 1.**
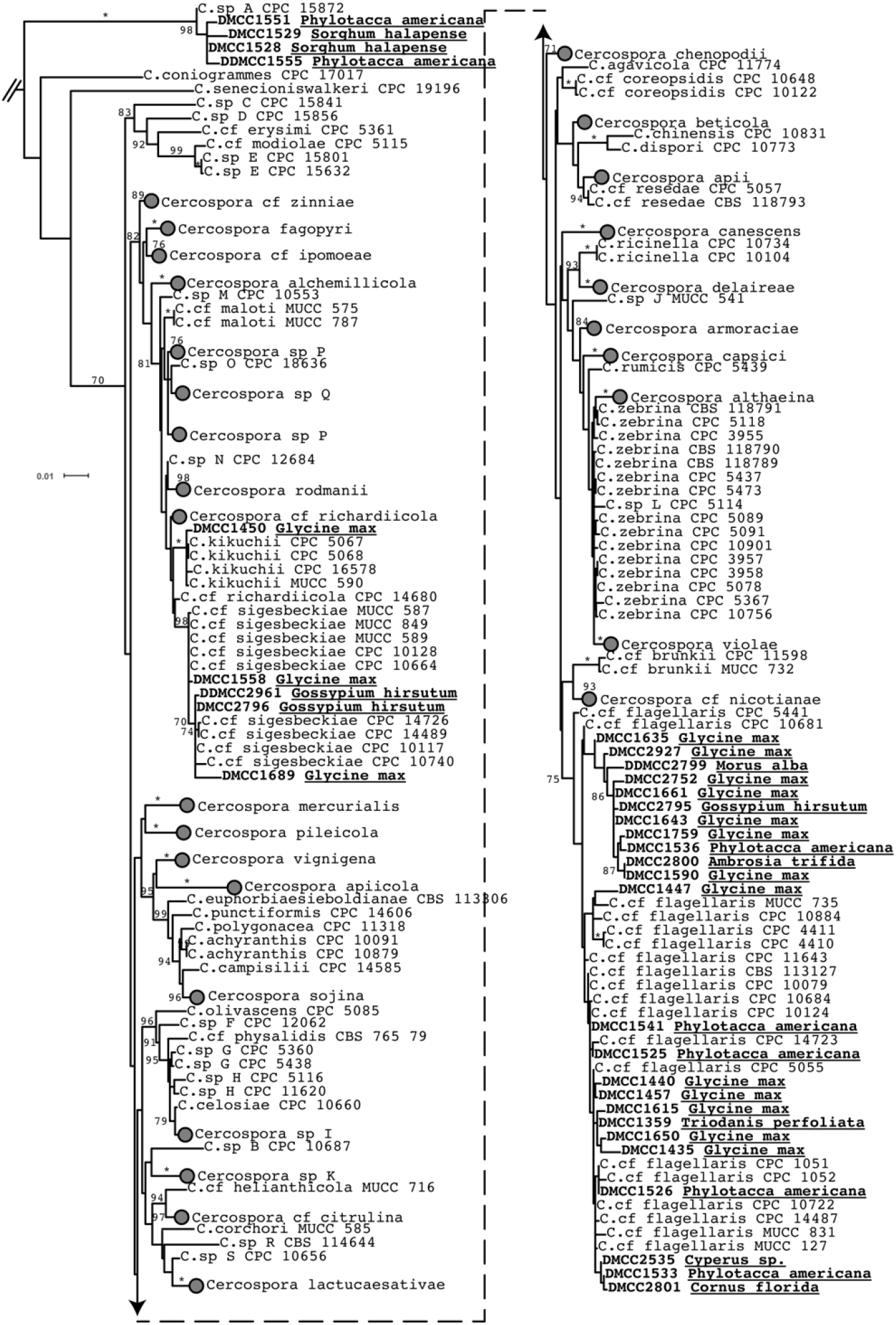
Maximum-likelihood phylogeny estimated from a concatenated alignment of actin, calmodulin, translation elongation factor 1α, histone 3, and nrITS. Bootstrap support values are above or adjacent to each node with support greater than 70%. Isolates in bold font were collected and sequenced for this study and the host plant of origin is underlined.

Mycelial tissue of *C*. cf. *flagellaris* isolate DMCC1359 was sent to SNPsaurus (Eugene, OR) for high-molecular weight DNA extraction, library preparation, and sequencing. A total of 6.14 Gb of PacBio CLR reads and 0.45 Gb of single-end Illumina sequencing reads were generated. Due to the inclusion of spike-in DNA during the PacBio CLR sequencing, we leveraged the Illumina short-read sequencing data to remove potential contaminants. Illumina sequences were assembled using Spades with automatic k-mer selection (Bankevich et al. 2012). PacBio reads were mapped to the short read assembly using minimap2 (Li 2018) and mapped reads were extracted using samtools (Li et al. 2009). The mapped PacBio reads were then independently assembled using Canu (Koren et al. 2017) and Flye (Kolmogorov et al. 2019) with default parameters. Contiguity of each assembly was assessed with Quast (Gurevich et al. 2013) and telomeric repeats (TTAGGG/CCCTAA) were identified using a python script (https://github.com/JanaSperschneider/FindTelomeres). We observed that the Flye assembly was slightly more contiguous, however, the Canu assembly contained more completely assembled subtelomeric regions (Table 1). To maximize contiguity while retaining well-assembled subtelomeric regions, we next scaffolded the two assemblies using Ragtag (Alonge et al. 2022), using the draft Flye assembly as the reference and the Canu assembly as the query. The Illumina short reads were then mapped to the scaffolded assembly using BWA-MEM (Li 2013) and the alignments were used for polishing using pilon (Walker et al. 2014). The polished scaffolded assembly was highly contiguous and consisted of 14 scaffolds, a total assembly size of 33.72 Mb, an N50 of 2.90 Mb, and an L50 of 5 (Table 1). Four scaffolds had telomeric repeats on both ends, signifying completely assembled chromosomes. Nine additional scaffolds had telomeric repeats on only one end, representing partially assembled chromosomes. Additionally, one 44.1 kb scaffold (scaffold_15), represented the mitochondrial genome. Genome assembly completeness was assessed using BUSCO (Simão et al. 2015) with the ascomycota_odb10 database, which revealed a 99.50% completeness. The raw sequencing data and scaffolded genome assembly was deposited in the NCBI database (BioProject PRJNA1466299; BioSample SAMN60044423).

**Table 1.** Genome assembly statistics.

| Parameter | Canu | Flye | Scaffolded Assembly |
| --- | --- | --- | --- |
| Assembly Size | 33.55 | 33.89 | 33.72 |
| Contigs | 29 | 17 | 15 |
| N50 | 2.04 | 2.80 | 2.90 |
| L50 | 7 | 6 | 5 |
| Telomeres | 19 | 4 | 17 |
| Telomere-to-telomere contigs | 2 | 0 | 4 |
| BUSCO<sup>1</sup> | 99.50% | 99.50% | 99.50% |
| Genes | NA | NA | 11,268 |
| Secreted Proteins | NA | NA | 947 |
| Effectors | NA | NA | 360 |
| CAZymes<sup>2</sup> | NA | NA | 512 |
| BGCs<sup>3</sup> | NA | NA | 60 |

<sup>1</sup>Complete benchmarking universal single copy orthologs identified in assembly using the ascomycota\_odb10 database
<sup>2</sup>Carbohydrate-active enzymes identified from the dbCAN3 server
<sup>3</sup>Biosynthetic gene clusters identified from AntiSmash (Fungal Version)

Next, we assessed repetitive element content of the scaffolded assembly. RepeatModeler and RepeatMasker (Flynn et al. 2020; Smit et al. 2013) were used to generate a custom repeat library and mask the assembly, respectively, revealing an overall 2.62% repeat content. The masked assembly was then annotated with the funannotate pipeline (Palmer and Stajich 2019) using the UniProt protein database as protein evidence and resulted in the annotation of 11,268 genes. The genome annotation is available in a public GitHub repository (https://github.com/jkzrich/Cercospora_flagellaris_genome). SignalP v5 (Almagro Armenteros et al. 2019) and EffectorP (Sperschneider and Dodds 2022) were then used to predict secreted proteins and effectors, respectively. A total of 947 genes were predicted to encode secreted proteins, of which, 360 were predicted to be effectors. We next annotated secondary metabolite gene clusters using AntiSmash (Blin et al. 2025) and identified 60 biosynthetic gene clusters. Finally, carbohydrate-active enzymes (CAZymes) were predicted using the dbCAN3 server (Zheng et al. 2023). Genes classified as CAZymes by at least two tools (HMMER:dbCAN, HMMER:dbCAN-sub, or DIAMOND:CAZy) were retained, resulting in the identification of 512 genes encoding CAZymes.

In summary, we assembled a highly contiguous reference genome for the economically important soybean pathogen *C*. cf. *flagellaris*. This genome assembly will be a valuable resource for future comparative, functional, and population genomics studies that will address fundamental questions in this economically important pathosystem.

